# Animal movement in pastoralist populations and implications for pathogen spread and control

**DOI:** 10.1101/2020.02.12.946467

**Authors:** George P. Omondi, Vincent Obanda, Kimberly VanderWaal, John Deen, Dominic A. Travis

## Abstract

Infectious diseases are one of the most important constraints to livestock agriculture, and hence food, nutritional and economic security in developing countries. In any livestock system, the movement of animals is key to production and sustainability. This is especially true in pastoralist systems where animal movement occurs for a myriad of social, ecological, economic and management reasons. Understanding the dynamics of livestock movement within an ecosystem is important for disease surveillance and control, yet there is limited data available on the dynamics of animal movement in such populations. The aim of this study was to investigate animal transfer networks in a pastoralist community in Kenya, and assess network-based strategies for disease control. We used network analysis to characterize five types of animal transfer networks and evaluated implications of these networks for disease control through quantifying topological changes in the network because of targeted or random removal of nodes. To construct these networks, data were collected using a standardized questionnaire (N=164 households) from communities living within the Maasai Mara Ecosystem in southwestern Kenya. The median livestock movement distance for agistment (dry season grazing) was 39.49 kilometers (22.03-63.49 km), while that for gift, bride price, buying and selling were 13.97 km (0-40.30 km), 30.75 km (10.02-66.03 km), 31.14 km (17.56-59.08 km), and 33.21 km (17.78-58.49 km), respectively. Our analyses show that the Maasai Mara National Reserve, a protected area, was critical for maintaining connectivity in the agistment network. In addition, villages closer to the Maasai Mara National Reserve were regularly used for dry season grazing. In terms of disease control, targeted removal of highly connected village nodes was more effective at fragmenting each network than random removal of nodes, indicating that network-based targeting of interventions such as vaccination could potentially disrupt transmission pathways and reduce pathogen circulation in the ecosystem. In conclusion, this work shows that animal movements have the potential to shape patterns of disease transmission and control in this ecosystem. Further, we show that targeted control is a more practical and efficient measure for disease control.

## Introduction

Rangeland ecosystems in Africa, defined as areas of natural or semi-natural vegetation in arid or semi-arid climates, host large numbers of wildlife, livestock, and marginalized pastoralist populations (Homewood, 2004; Le Houerou, 2012). Low rainfall and seasonally heterogeneous resources characterize such areas, which necessitates human and livestock mobility to utilize spatiotemporally distributed resources (Swallow, 1994; Butt, 2010; Goldman and Riosmena, 2013; Turner and Schlecht, 2019). However, animal movements have been shown to impact disease patterns (Fevre et al., 2006; Altizer et al., 2011), especially among pastoral communities (Rajeev et al., 2017; Sintayehu et al., 2017; VanderWaal et al., 2017), where losses emanating from livestock diseases affect livelihoods, and their control has the potential to enhance household productivity and health outcomes (Marsh et al., 2016). Rangeland systems are especially at high risk for pathogen introduction and spread because grazing livestock interact with both wildlife and other livestock directly and indirectly through shared forage and water resources (Rajeev et al., 2017). Thus, control of infectious diseases in livestock systems requires an understanding of interaction not only between herds but also in different locations (Bastos et al., 2000; Fevre et al., 2006; zu Dohna et al., 2014; Machado et al., 2019; Omondi et al., 2019).

Patterns of contact between livestock herds influences the spread of infectious pathogens (Fevre et al., 2006) and thus can be used to characterize epidemiological dynamics (Kao et al., 2006; VanderWaal et al., 2016; VanderWaal et al., 2017) and develop targeted surveillance and control strategies (Bajardi et al., 2012; Frossling et al., 2014; Ribeiro-Lima et al., 2015). Herds with high rates of between-herd contacts have a higher risk of acquiring infections (VanderWaal et al., 2017). In addition, infections often propagate from a small number of actors (Woolhouse et al., 1997; Volkova et al., 2010), with so-called “super-spreaders” disproportionately contributing to transmission events (Lloyd-Smith et al., 2005). Thus, characterizing the underlying architecture of contact patterns within a population can help elucidate important drivers and pathways for disease transmission as well as critical points for surveillance and control (Bajardi et al., 2012; VanderWaal et al., 2016).

Pastoralists have adopted a strategy of complex livestock movement to maximize the utilization of seasonally available resources, leading to complex heterogeneous contact patterns and variability in disease outcomes (Dejene et al., 2016; Rajeev et al., 2017; Sintayehu et al., 2017; Turner and Schlecht, 2019). In such communities, livestock movement data is seldom available, and thus contact is difficult to characterize. Several studies have attempted to model livestock movement by analyzing sales records (Chaters et al., 2019), animal transaction records combined with questionnaire surveys (Motta et al., 2017), census of migrating pastoralists (Pomeroy et al., 2019), Global Positioning System data loggers (VanderWaal et al., 2017), and ego-based approaches (Bronsvoort et al., 2004). However, none of the methods above captures the diversity of social drivers behind movements within pastoral cultural systems. For instance, in addition to buying and selling, Maasai pastoralists move animals, with or without the transfer of ownership, for instance for bride price payments, lending of animals between friends and families, gifts, and for seasonal access to pasture and water (Perlov, 1987 in (Aktipis et al., 2016). In pastoralist populations, moving or sharing animals is both a survival strategy, a relationship building exercise, and often a method of risk pooling (Aktipis et al., 2011; Aktipis et al., 2016). Pastoralists communities are known to use gifts of livestock as a means to build and enhance relationships (de Vries et al., 2006). The role that such livestock movements play in disease dissemination is seldom evaluated, but may be key to maximizing productivity of this management system (Sintayehu et al., 2017).

Designing control strategies is complex, and traditional epidemiological approaches often fail to capture the dynamic, non-linear, and interconnected nature of pastoral systems (Benham-Hutchins and Clancy, 2010). To further our understanding of cattle-associated movement dynamics, graph theory can be used to quantify within- and- between village movements such that household actions, for instance buying/selling, connect different villages. Such between-village connections are expected to be important when developing disease surveillance and intervention strategies (Watts, 1987; Woolhouse et al., 1997; Sintayehu et al., 2017; Ahmed et al., 2018; Russell et al., 2018). In graph theory, networks are used to characterize interacting systems in which nodes (here defined as households or villages) are inter-connected through edges (here defined as movements (Craft and Caillaud, 2011; Danon et al., 2011; Silk et al., 2017; Sintayehu et al., 2017; Balasubramaniam et al., 2018; Ogola et al., 2018). In our study, a network edge is a potential route for transmission of pathogens between households through the movement of animals. Using network analysis, we can calculate centrality metrics to evaluate the importance of a node in connecting the network, investigate the propagation of a hypothetical disease, and assess the potential for targeted surveillance or control if a node is removed - a measure equivalent to vaccination or depopulation (Martinez-Lopez et al., 2009; Kinsley et al., 2019; Yang et al., 2019). In this study, our objective was to use network analysis to characterize different types of animal movement, and evaluate their potential role in disease transmission and control in a pastoralist community in Kenya. We hypothesized that villages proximal to Maasai Mara National Reserve will play an important role in the connectivity of the ecosystem, as measured by their centrality metrics, and that targeted control measures aimed at villages with the most connections will be more efficient at fragmenting the connectivity of the network than a non-targeted approach. This study advances our understanding of the movement dynamics of livestock within a pastoralist community characterized by variable animal transfer pathways and their role in network-based interventions for livestock disease surveillance and control.

## Material and Methods

### Study site

This study examined the dynamics of livestock movement in pastoralist communities living within the Maasai Mara Ecosystem (MME) (Figure 1). MME is located in southwestern Kenya, and encompasses the 1,530 km^2^ MMNR, within which livestock grazing is banned and adjoining pastoral ranches where communal settlements, livestock grazing, and tourism are permitted (Bhola et al., 2012). The livestock movement dynamics of the communities in this ecosystem are driven by the Maasai Mara National Reserve (MMNR) due to the availability of forage within this wildlife conservation area during the dry season (Reid et al., 2003; Butt et al., 2009). Rainfall in this ecosystem is largely bimodal, varying from 500 mm in the southeast to 1300 mm in the northwest (Bartzke et al., 2018). These factors combine to create spatiotemporal heterogeneity in water and forage distribution, which influences wild herbivores and domestic stock movement within the ecosystem. This ecosystem is located within the larger Narok County, which is a 17,953 km^2^ area with more than one million cattle, 2.3 million sheep and goats, and a human population that is largely rural (KNBS, 2010).

**Figure 1:**
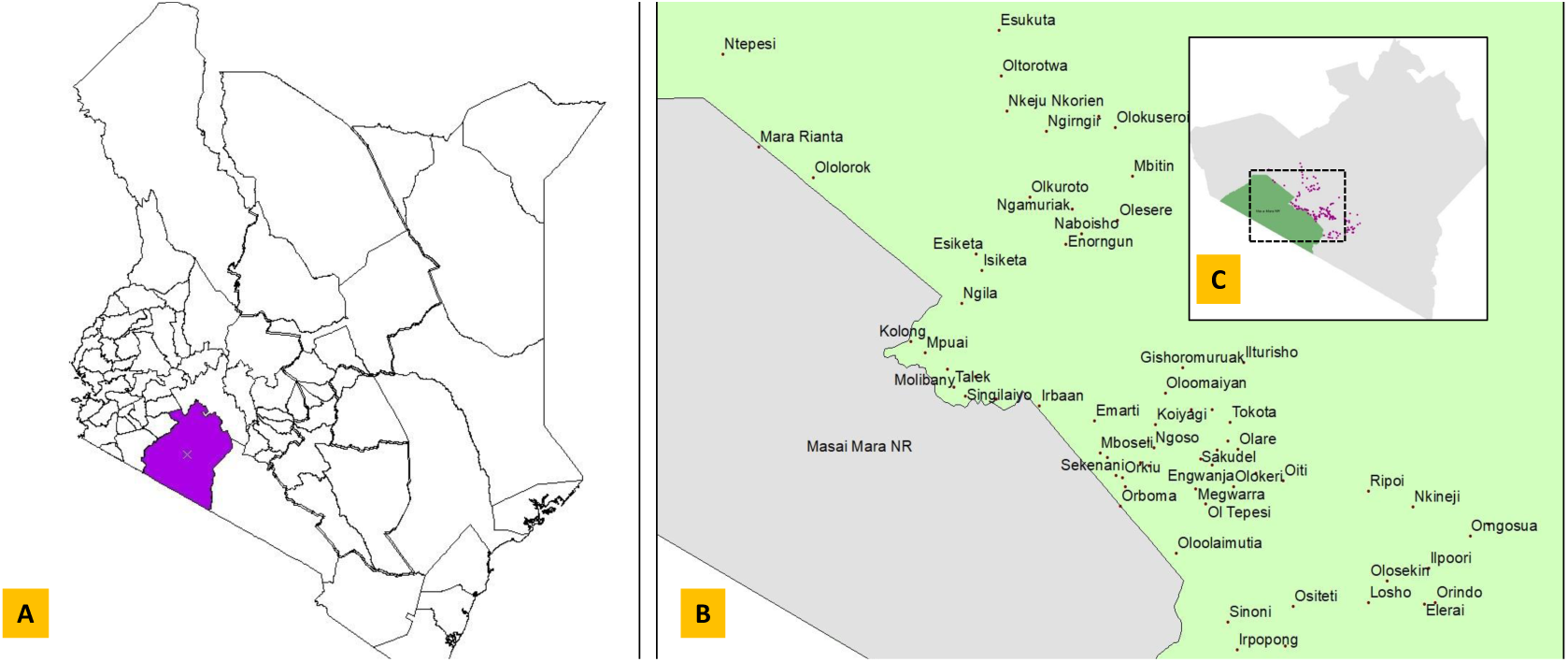
Map of the Maasai Mara Ecosystem. A. Map of Kenya with Narok County in red; B. Unique villages sampled in this study; C. Map of Narok County with households sampled marked in purple, dotted square represents an area equivalent to B.

### Data collection

This research is part of a larger study aimed at understanding zoonotic disease occurrence in the Maasai Mara Ecosystem, with all sampling conducted between November 2017 and June 2019. We defined households as persons living within an abode for a period of one month prior to the sampling, and herds as groups of cattle, sheep and goats, and any other domestic stock owned by the respondent. We purposively sampled one hundred and sixty four households, targeting those within 20 kilometers of the Maasai Mara National Reserve. Pastoral cattle tend to move longer daily distances than small stock, with an average of 2-9 kilometers being the norm for grazing (Turner and Schlecht, 2019). For longer-term migration, or “travel mobility,” in pastoralist systems, the average distance moved ranges from 47-170 kilometers (Turner and Schlecht, 2019), with the wide variation indicative of an individual household’s cost-benefit valuation of the move.

For this study, we defined five animal transfer pathways identified from an initial scoping survey in the ecosystem; agistment (defined as the temporary re-location of animals to access forage and water in other locations during the dry season, usually lasting 2-3 months, while maintaining a household in a single village), gift, bride price, buying, and selling. We interviewed a household respondent using a structured questionnaire, in which the respondent was asked to identify villages (by common name) from which they either sent or received animals through any of the aforementioned pathways over the last five years. The respondents were requested to name villages rather than specific household due to the logistical constraints of collecting locational data on households named by respondents. For each household interviewed and village named, locational data was recorded using a handheld Global Positioning System; a centroid was calculated to represent the location of villages in which multiple household locations were recorded. The University of Minnesota IRB (STUDY00000837), Kenya Wildlife Service (KWS/BRM/5001), and the Narok County government (NCG/HEALTH/GEN/VOL.1/2) authorized this study.

### Data analysis

#### Network construction

In graph theory, nodes can be partitioned into *k* independent sets or groups. A network with *k* ≥ 3 is a *multipartite* network, whereas those with one or two independent sets or groups are *unipartite* and *bipartite* networks, respectively (Jacoby and Freeman, 2016). We constructed a household-village bipartite network, where households are linked to villages to which they received or moved animals. In this study, a separate bipartite network was constructed for each type of contact. In a bipartite network B = {*U, V, E*}, where U and V are the disjoint set of nodes representing households and villages, respectively, and E is the linkage between nodes, such as E = {(*u, v*): u ∈ *U, v* ∈ *V*}. In this network, nodes in *U* can only connect to nodes in *V*, and no connections among nodes of the same type exist (Banerjee et al., 2017). This representation of contact is appropriate for the method by which data were collected for this project (households were asked about movements of animals to different villages). These data can be represented by an unweighted biadjacency matrix B= {*U, V, E*}, which is a (0, 1) matrix of size |*U*|×|*V*|; B_*uv*_ =1 if there is an edge between *u* and *v*, or B_*uv*_ = 0 when there is none. Thus, households are connected to other households indirectly based on villages to which they had common connections. In this sense, each set of nodes (villages and households) have independent properties that we can estimate to evaluate the roles played by each set. These properties will be evaluated at two levels, first, a household’s role within the network, and secondly the villages’ role in the network.

#### Network metrics

At the node-level, we calculated two centrality metrics: degree and betweenness. We also summarized the density and fragmentation index of the network as a whole (Table 1), and visualized network topology. All analysis were conducted using the *igraph* package (Csardi, 2013).

**Table 1:** Definition of network metrics

| Node-level Metrics |  | Citation |
| --- | --- | --- |
| Degree | Total number of unique nodes that sent or received an animal to/from a particular node. | (Wasserman and Faust, 1994) |
| Betweenness | Number of times a node is located on the shortest path between any two pairs of nodes within the observed network | (Wasserman and Faust, 1994) |
| Network-level Metrics |  |  |
| Density | Number of observed contacts in the network relative to all possible contacts. | (Wasserman and Faust, 1994) |
| Fragmentation index | Proportion of pairs of nodes that are disconnected (no paths exist connecting them) in the network. | (Chen et al., 2007) |

#### Implications of node removal

We used two approaches for node removal, random and targeted. In random removal, we selected any 2, 5, or 10 nodes at random, calculated network-level metrics before and after removal, and repeated this process for 1,000 iterations to generate an expected distribution. For targeted removal, we selected top 2, 5, and 10 nodes based on degree, and recalculated the network-level metrics before and after removal (Albert et al., 2000; Holme et al., 2002; Chen et al., 2007). We quantified the topological impact of removing nodes using the fragmentation index, *F*, which is the proportion of non-connected pairs of nodes in the network. *F* = 0 would represent a fully-connected, non-fragmented network in which all pairs of nodes are connected through paths in the network, and *F* = 1 would represent a fully fragmented network where every node is isolated (Borgatti, 2006; Chen et al., 2007).

## Results

We sampled 164 households in 67 unique villages (Figure 1 B) across the Maasai Mara Ecosystem, with 30% of the respondents being female (50/164), 70% male (114/164), and the median length of time they had lived in the area being 8 years (4-100 years). Of the respondents interviewed, 99% identified as pastoralists, though 12% also reported formal employment, and 4% were merchants involved in various trades.

### Network metrics and visualization

Five animal movement ‘reasons’ were examined; agistment, bride price, gift, buying, selling. The median livestock movement distance for agistment was 39.49 kilometers (22.03-63.49 km), while that for gift, bride price, buying, and selling were 13.97 km (0-40.30 km), 30.75 km (10.02-66.03 km), 31.14 km (17.56-59.08 km), and 33.21 km (17.78-58.49 km), respectively. For agistment, gift, bride price, buying and selling networks, network densities were 0.0038, 0.0023, 0.0022, 0.0082, and 0.0056, respectively. In addition, we summarized the degree of the villages and the households in our networks separately. The median household degrees in agistment, gift, bride price, buying and selling network node degrees were 2 (interquartile range: 1-2), 1 (0-1), 1 (1-2), 2 (1-3), and 2 (1-3), respectively. We report the median degree for households only, as the interpretation of degree for villages is less straight forward because some villages were identified by a respondent but not sampled during questionnaire interviews. With that caveat, seven villages were shown to be important across all networks. These included Sekenani, Talek, Ololaimutia, Nkineji, Olesere, Nkoilale and Naikara (Table 2). Of the villages evaluated, Maasai Mara National Reserve (though technically not a village) had the highest degree and betweenness for agistment (Table 2).

**Table 2:**
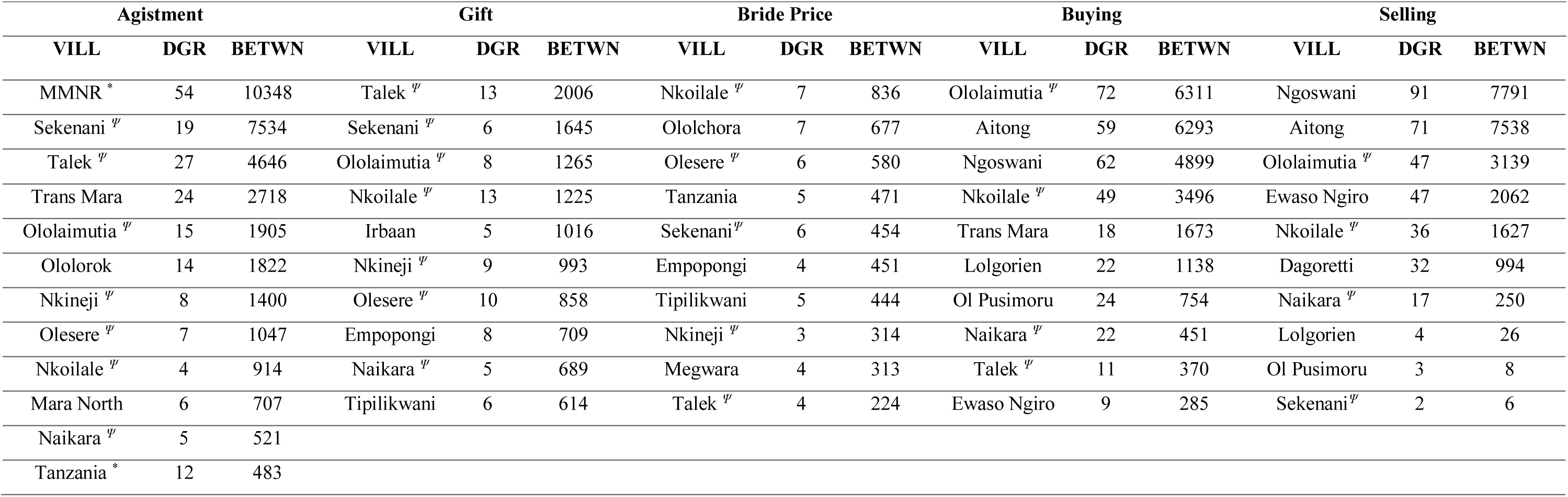
Network metrics; degree and betweenness of the top 10 villages in the five networks evaluated in this study. * Not a village within the ecosystem but serves as an important hub in the network – this was only in the agistment network. ^Ψ^ Villages that are common across all evaluated networks. VILL = Village; DGR = degree; BETWN = betweeness.

| Agistment |  |  | Gift |  |  | Bride Price |  |  | Buying |  |  | Selling |  |  |
| --- | --- | --- | --- | --- | --- | --- | --- | --- | --- | --- | --- | --- | --- | --- |
| VILL | DGR | BETWN | VILL | DGR | BETWN | VILL | DGR | BETWN | VILL | DGR | BETWN | VILL | DGR | BETWN |
| MMNR * | 54 | 10348 | Talek <sup>ψ</sup> | 13 | 2006 | Nkoilale <sup>ψ</sup> | 7 | 836 | Ololaimutia <sup>ψ</sup> | 72 | 6311 | Ngoswani | 91 | 7791 |
| Sekenani <sup>ψ</sup> | 19 | 7534 | Sekenani <sup>ψ</sup> | 6 | 1645 | Ololchora | 7 | 677 | Aitong | 59 | 6293 | Aitong | 71 | 7538 |
| Talek <sup>ψ</sup> | 27 | 4646 | Ololaimutia <sup>ψ</sup> | 8 | 1265 | Olesere <sup>ψ</sup> | 6 | 580 | Ngoswani | 62 | 4899 | Ololaimutia <sup>ψ</sup> | 47 | 3139 |
| Trans Mara | 24 | 2718 | Nkoilale <sup>ψ</sup> | 13 | 1225 | Tanzania | 5 | 471 | Nkoilale <sup>ψ</sup> | 49 | 3496 | Ewaso Ngiro | 47 | 2062 |
| Ololaimutia <sup>ψ</sup> | 15 | 1905 | Irbaan | 5 | 1016 | Sekenani <sup>ψ</sup> | 6 | 454 | Trans Mara | 18 | 1673 | Nkoilale <sup>ψ</sup> | 36 | 1627 |
| Ololorok | 14 | 1822 | Nkineji <sup>ψ</sup> | 9 | 993 | Empopongi | 4 | 451 | Lolgorien | 22 | 1138 | Dagoretti | 32 | 994 |
| Nkineji <sup>ψ</sup> | 8 | 1400 | Olesere <sup>ψ</sup> | 10 | 858 | Tipilikwani | 5 | 444 | Ol Pusimoru | 24 | 754 | Naikara <sup>ψ</sup> | 17 | 250 |
| Olesere <sup>ψ</sup> | 7 | 1047 | Empopongi | 8 | 709 | Nkineji <sup>ψ</sup> | 3 | 314 | Naikara <sup>ψ</sup> | 22 | 451 | Lolgorien | 4 | 26 |
| Nkoilale <sup>ψ</sup> | 4 | 914 | Naikara <sup>ψ</sup> | 5 | 689 | Megwara | 4 | 313 | Talek <sup>ψ</sup> | 11 | 370 | Ol Pusimoru | 3 | 8 |
| Mara North | 6 | 707 | Tipilikwani | 6 | 614 | Talek <sup>ψ</sup> | 4 | 224 | Ewaso Ngiro | 9 | 285 | Sekenani <sup>ψ</sup> | 2 | 6 |
| Naikara <sup>ψ</sup> | 5 | 521 |  |  |  |  |  |  |  |  |  |  |  |  |
| Tanzania * | 12 | 483 |  |  |  |  |  |  |  |  |  |  |  |  |

**Figure 2:**
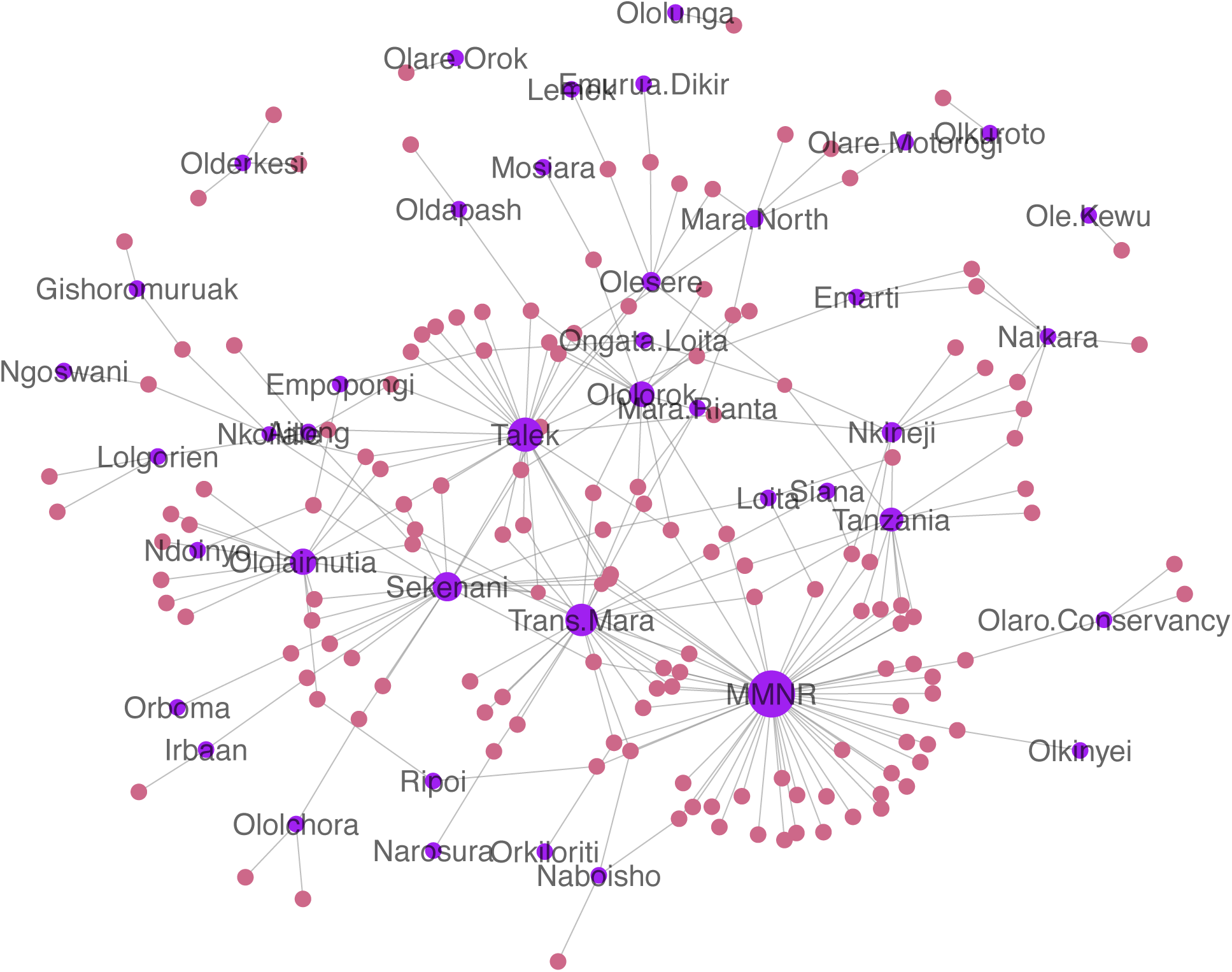
Agistment Network: Bipartite network of agistment locations within the Maasai Mara ecosystem, with node sizes scaled by degree. Purple nodes are villages, while pale violet nodes are households.

**Figure 3:**
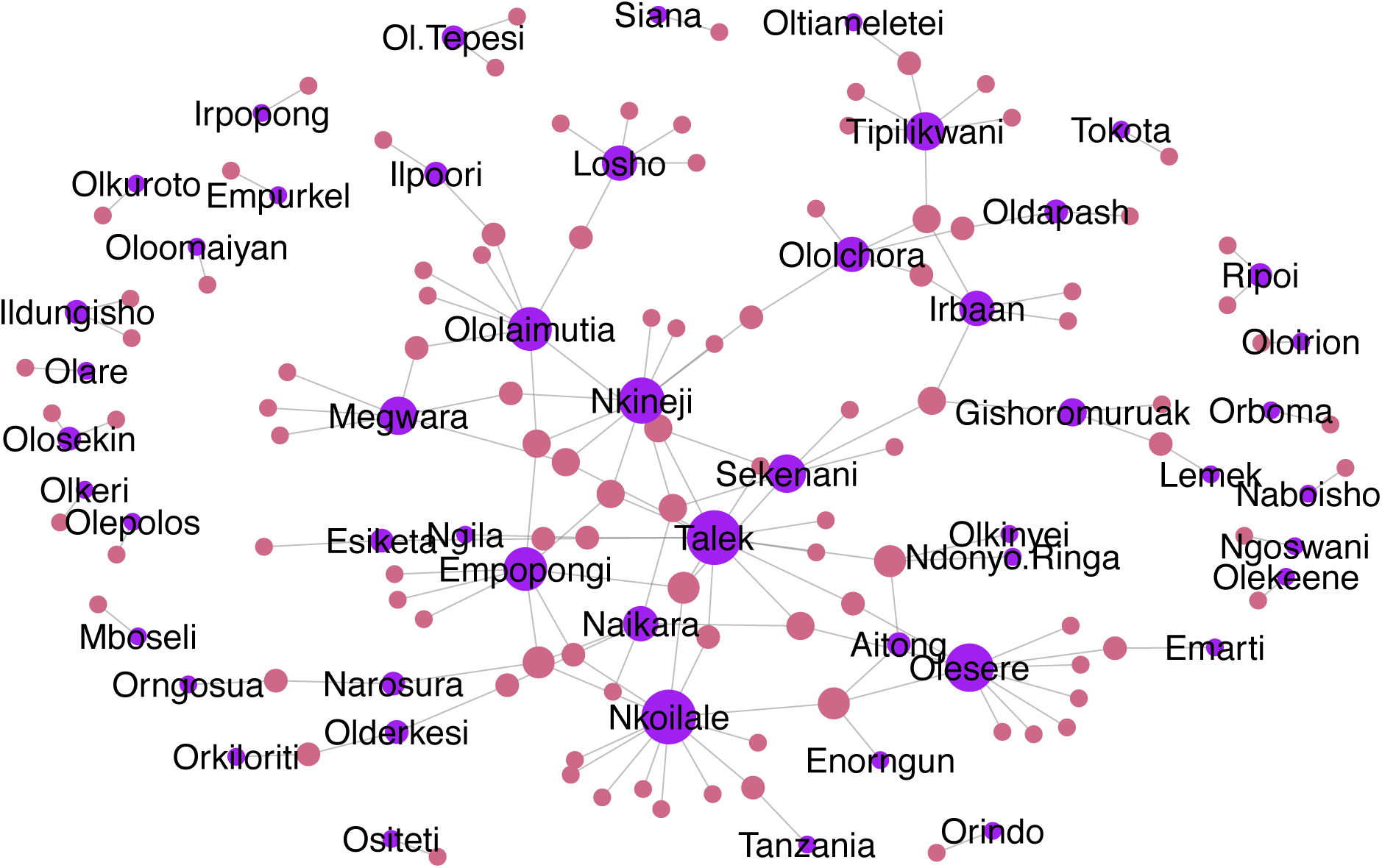
Gift Network: Bipartite network of gift locations within the Maasai Mara ecosystem, with node sizes scaled by degree. Purple nodes are villages, while pale violet nodes are households.

**Figure 4:**
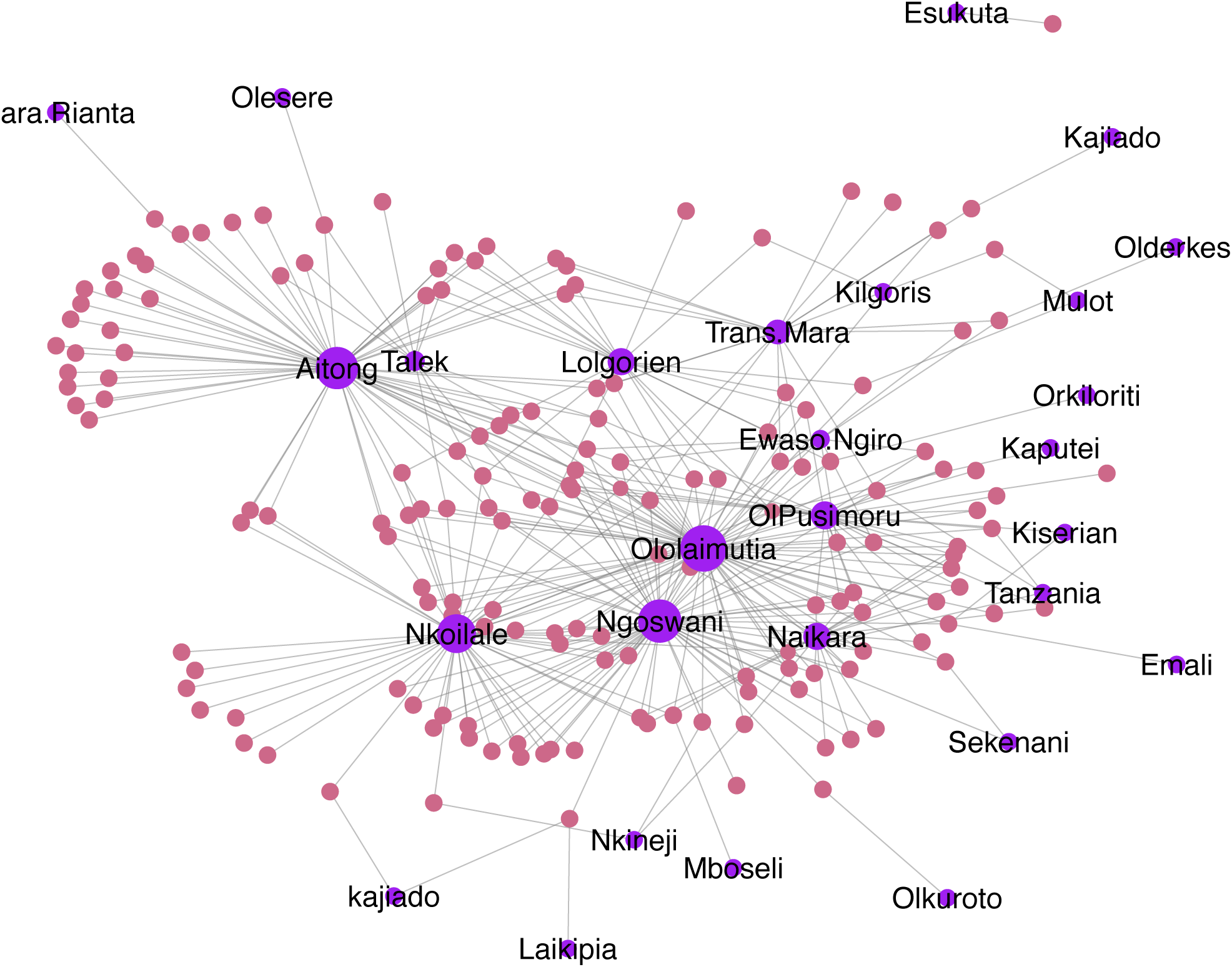
Buying Network: Bipartite network of major buying locations within the Maasai Mara ecosystem, with node sizes scaled by degree. Purple nodes are villages, while pale violet nodes are households.

### Implications of node removal

We used the fragmentation index to examine changes in network topology following the targeted and random removal of nodes in the five networks. Consistent with our hypothesis, targeted removal outperformed random removal of nodes in terms of increasing network fragmentation. The fragmentation indices of the full networks were 0.51, 0.83, 0.93, 0.45 and 0.47 for agistment, gift, bride price, buying and selling, respectively. Across all network types, targeted removal of nodes based on their degree resulted in substantially higher fragmentation than random removal of nodes; the fragmentation indices for the targeted removals always exceeded the upper bounds of the 95% interval that was achieved through random removals. This result was consistent regardless of whether the top 2, 5, or 10 nodes with highest degree were removed (Table 2). The biggest change in fragmentation was realized when five nodes were targeted for removal, with only a modest additional benefit in removing 10 nodes as opposed to five (complete list of figures are included in the supplementary materials).

**Figure 5:**
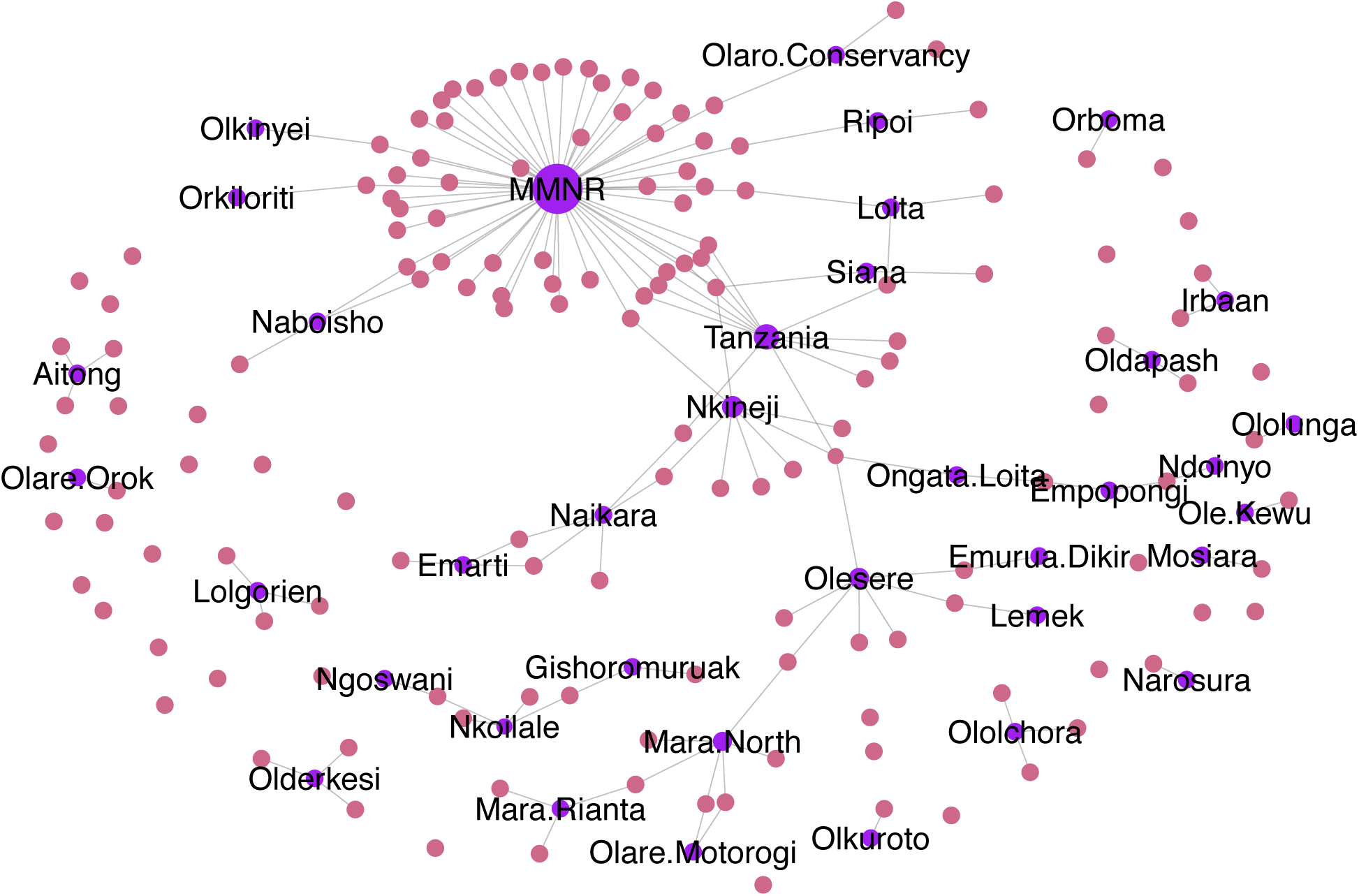
Bipartite network of agistment locations following targeted removal of top five villages. Purple nodes are villages, while pale violet nodes are households.

**Table 3:** Network metrics; fragmentation index of the five networks evaluated in this study following the removal of 2, 5 and 10 nodes, with nodes removed either selected randomly or targeted based on degree. For random removals, the median (95% confidence interval) fragmentation is reported, summarized across 1,000 iterations.

| Fragmentation of the network following targeted versus random node removal |  |  |  |  |  |  |  |
| --- | --- | --- | --- | --- | --- | --- | --- |
| Reason | Full Network | Removal of 2 villages |  | Removal of 5 villages |  | Removal of 10 villages |  |
|  | Targeted | Random | Targeted | Random | Targeted | Random | Targeted |
| Buying | 0.45 | 0.54 | 0.48 (0.45-0.59) | 0.86 | 0.53 (0.47-0.67) | 1.00 | 0.64 (0.51-0.82) |
| Selling | 0.47 | 0.71 | 0.49 (0.47-0.66) | 0.94 | 0.56 (0.49-0.73) | 1.00 | 0.75 (0.52-0.89) |
| Agistment | 0.51 | 0.61 | 0.54 (0.52-0.59) | 0.81 | 0.58 (0.53-0.65) | 0.90 | 0.64 (0.57-0.74) |
| Gift | 0.83 | 0.89 | 0.84 (0.83-0.87) | 0.96 | 0.86 (0.83-0.91) | 0.99 | 0.89 (0.85-0.94) |
| Bride Price | 0.93 | 0.97 | 0.94 (0.93-0.96) | 0.99 | 0.95 (0.94-0.98) | 1.00 | 0.97 (0.95-0.99) |

## Discussion

In this study, our goal was to characterize animal transfer networks in pastoralist communities in Kenya, and evaluate their potential role in disease control and management. As described in Table 1, degree, betweenness, density, and fragmentation index are important measures of a network’s topology and for assessing potential impact of perturbing this topology for disease control (Wasserman and Faust, 1994; Chen et al., 2007). We showed that animals were moved the longest distances (median = 39.49 km) for agistment (movement of animals to forage and water during the dry season), followed by bride price, buying and selling, which were approximately similar (ranging between 30-33 km). Finally, movement due to gifting was more localized (with the lowest median distance at 13.97 km). Thus this study shows that buying, selling and agistment driven movements potentially play a bigger role than gift and bride price in disease propagation risk (with respect to both higher network densities and longer distance of movement). The protected area, Maasai Mara National Reserve, played an important role in connecting the ecosystem in that it was highly used for dry season grazing, as shown by its highest degree and betweenness. In addition, our results support the hypothesis that villages proximal to Maasai Mara National Reserve (Sekenani, Talek, and Ololaimutia) were more connected in the ecosystem (highest centrality metrics). In all networks, targeted removal of villages served to better fragment the network than randomly removing nodes, highlighting the potential benefits of targeted disease control strategies. Thus, targeted removal, such as vaccination, may provide an efficient approach for disease control in the ecosystem.

The fact that villages closer to MMNR were used regularly for dry season grazing is not a surprise given that, although grazing in the reserve is banned, it has been reported previously in the literature (Ogutu et al., 2009). In this study, respondents consistently identified the Maasai Mara National Reserve as a reservoir for forage during the dry season, even in the face of animal confiscation and fines levied by the County government (personal communication). The MMNR has become especially attractive following increased fencing of the ecosystem that has disrupted traditional animal foraging routes and grazing lands (Løvschal et al., 2017). Further, though buying and selling median distances were identical, a closer look at the network topology reveals more context. First, buying commonly includes transactions with villages outside Narok County (the study area). For instance, the respondents indicated that they bought their cattle from Tanzania, Kajiado, Kiserian, Emali, and Laikipia, all of which fall outside the county boundaries. This could be a strategy to acquire different or “better” livestock genetics (Ilatsia et al., 2012). On the contrary, selling of livestock mostly occurred in local markets. These included major markets such as Aitong, Nkoilale, Ololaimutia, Ewaso Ngiro, and Ol Pusimoru. In addition, a few farmers sold livestock in larger, peri-urban markets (e.g. Dagoretti, Ngong and Ongata Rongai) serving the capital city of Nairobi, possibly as a means of getting higher returns (Alarcon et al., 2017).

Network-based disease control relies on identifying a population’s contact structure and evaluating the role of the different nodes (e.g. villages, households, or farms) that could influence connectivity thus fragmenting the transmission network (Kiss et al., 2005; Tanaka et al., 2014). To evaluate the efficiency of network-based control strategies, we compared the effect of random versus targeted removal of nodes on the networks’ topological structure using the fragmentation index. Random removal of nodes requires no prior information on the network structure, but has been shown to be an inefficient approach (Albert et al., 2000). In our study, targeted removal of village nodes outperformed random removal, demonstrating the utility of network analysis in identifying highly connected villages that could be used for more strategic disease control or surveillance. Here, node removal mimics vaccination or depopulation, depending on the disease and context of infectious disease control (Keeling and Eames, 2005; Bansal et al., 2010). Ideally, an efficient fragmentation strategy should be one that removes minimal number of nodes as it represents, for instance, the minimum number of villages to be vaccinated to prevent further spread of an infection (Chen et al., 2007). We demonstrated that the removal of the top five nodes with the highest degree was effective at fragmenting all the networks. The agistment network, however, was more robust to node removal in that the removal of top 5 or 10 villages resulted in fragmentation indices of 81% and 90%, whereas this value was close to 100% for the other networks in this study. This might be due to the fact that, we cannot remove MMNR from the network or that household decisions to move to a particular location is highly influenced by an individual household’s cost-benefit analysis of the move independent of other household decisions (Turner and Schlecht, 2019). This is unlike buying and selling, which follow the law of supply and demand, and sometimes are dictated by intermediaries (Alemayehu, 2011; Alarcon et al., 2017; Chaters et al., 2019).

Agistment, buying, and selling networks occur much more frequently with potentially greater implications for pathogen dissemination than gifting and bride price (Macpherson, 1995; Bett et al., 2009). Anecdotally, we may conclude that the fragmentation of the selling network may serve to protect markets outside the Mara Ecosystem, such as Dagoretti, Ngong and Ongata Rongai. On the contrary, the fragmentation of the buying network leaves the markets in the neighboring counties connected to those in the Maasai Mara Ecosystem (Supplementary material) and may need to be considered when designing a comprehensive disease control strategy.

Our study has several limitations. First, data were collected at a single time point, and temporal changes in a network’s topology is a common phenomenon, especially in pastoralist production systems (VanderWaal et al., 2017; Pomeroy et al., 2019). Secondly, respondents were asked about movements made during the last five years, which limits the temporal resolution of when movements occurred and introduces potential recall bias. Third, because data were collected in a defined geographical area, the results may not be readily generalizable to other areas. Finally, our network structure did not account for common areas of daily contact, such as congregation during daily herding and at water resources, which may be important for localized disease transmission. Thus, our networks may under-represent connectivity amongst villages, particularly at local scales. However, our networks do represent longer distance movements in the ecosystem, with corresponding implications for longer distance pathogen spread.

## Conclusions

We have shown that the identification of highly connected villages could be beneficial in designing disease control programs that fragment potential transmission pathways in the livestock population. This fragmentation can be achieved through immunization of a node (node removal). Our findings demonstrate that even at a restricted spatial scale, network centrality measures may provide sufficient information to fragment networks, thus showing their utility not only for disease control but also in developing targeted risk-based surveillance approaches. Our approach of identifying villages rather than households has multiple advantages including cost implications and protection of privacy. However, the use of bipartite networks also allows for the identification of household nodes that may be relevant in the connecting the ecosystem. There is need however, to incorporate disease data from households in the ecosystem and evaluate the network topologies with respect to real-world transmission dynamics. In addition, it may be useful to consider economic costs of the information gathering and integrate risk analysis as a way to enhance the utility and robustness of the realized networks as presented here.

## Acknowledgements

We sincerely want to thank Samuel Soit, Joel Dukuny and James Lempoiyo for their immense help during the fieldwork portion of this research. In addition, we would like to thank the communities in the Maasai Mara Ecosystem for the patience and time in answering the questions and inquiries from the lead author. We would also like to thank the Kenya Wildlife Staff, specifically; Vasco Nyaga and Stephen Ndambuki Mwiu, for their immense support to the lead author in understanding the geography of the study area, and in addition extend our immense gratitude to all the Narok County members of staff for providing linkages to the communities in the Maasai Mara Ecosystem.

