## Supplementary Material for "Animal movement in pastoralist populations and implications for pathogen spread and control"

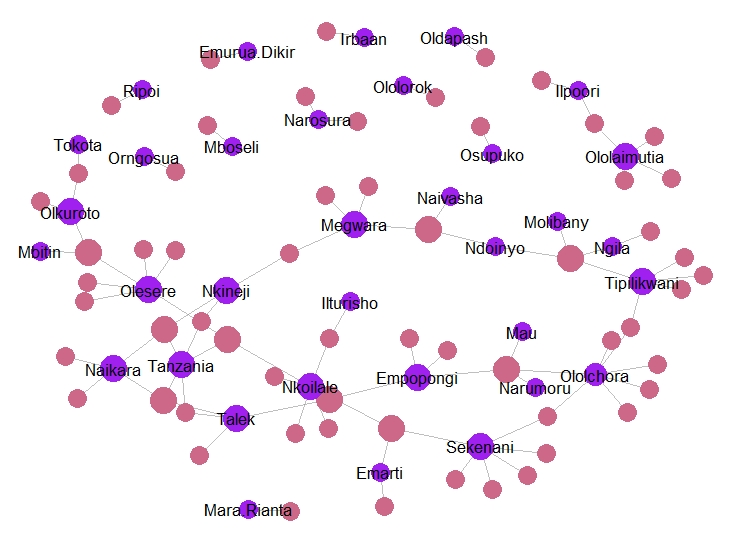
**Figure 1:** Bride Price Network: Bipartite network of bride price locations within the Maasai Mara ecosystem with node sizes scaled by degree. Purple nodes are villages, while pale violet nodes are households.

**Figure 2:** Selling Network: Bipartite network of major selling locations within the Maasai Mara ecosystem, with node sizes scaled by degree. Purple nodes are villages, while pale violet nodes are households.

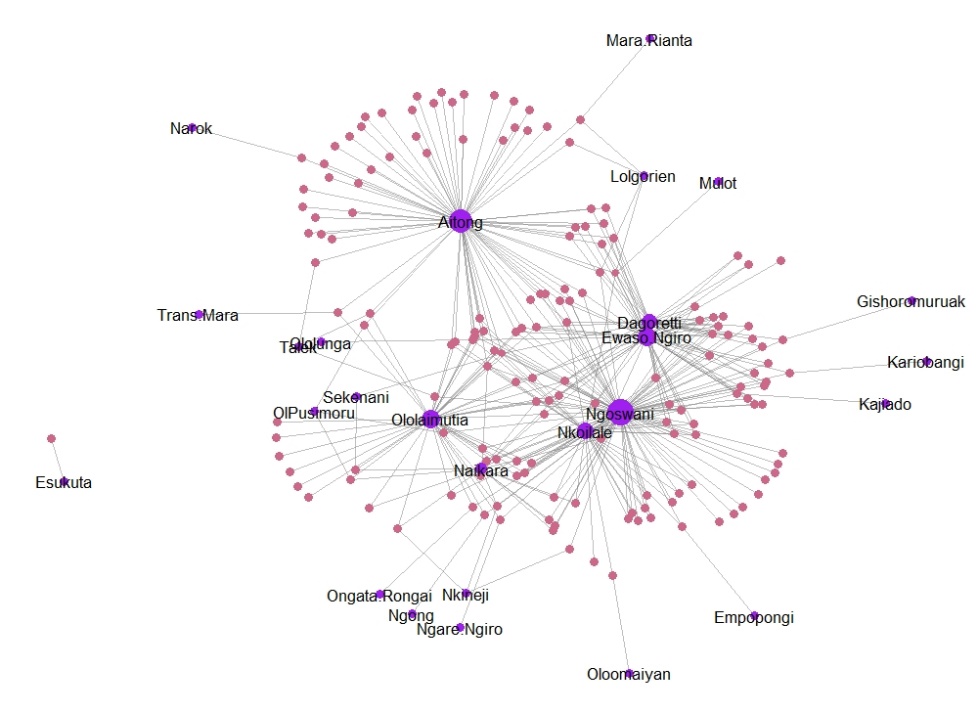

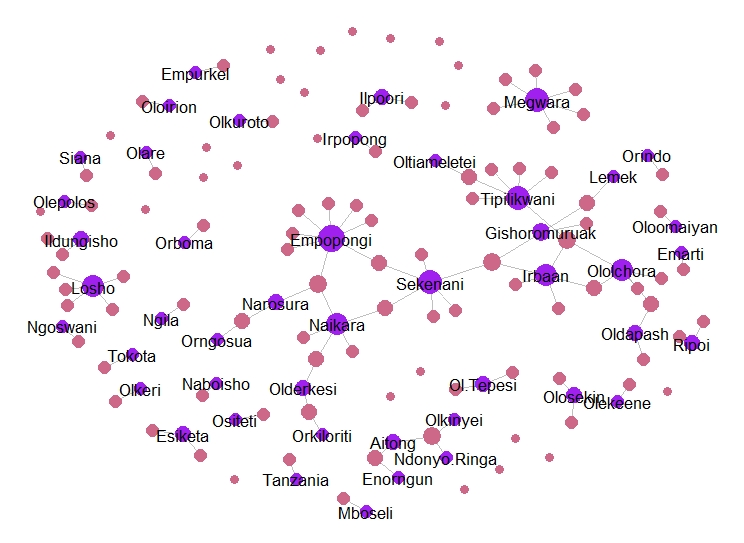
**Figure 3:** Bipartite network of gift locations following targeted removal of top five villages. Purple nodes are villages, while pale violet nodes are households.

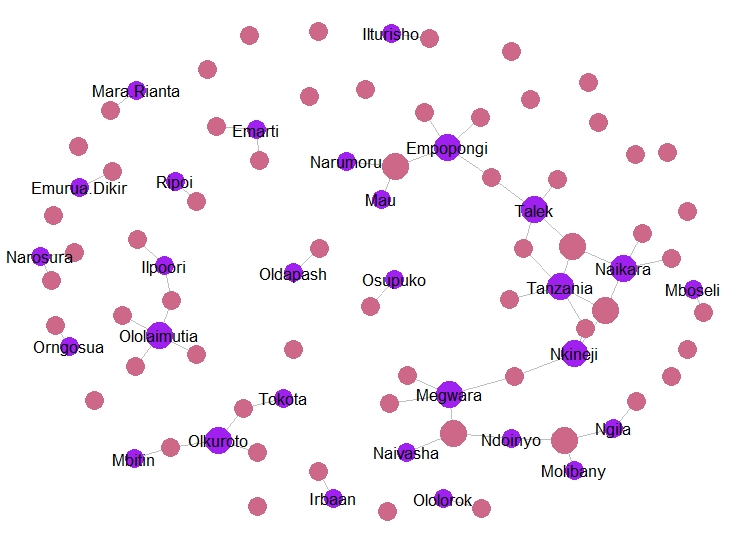
**Figure 4:** Bipartite network of bride price locations following targeted removal of top five villages. Purple nodes are villages, while pale violet nodes are households.

**Figure 5:** Bipartite network of buying locations following targeted removal of top five villages. Purple nodes are villages, while pale violet nodes are households.

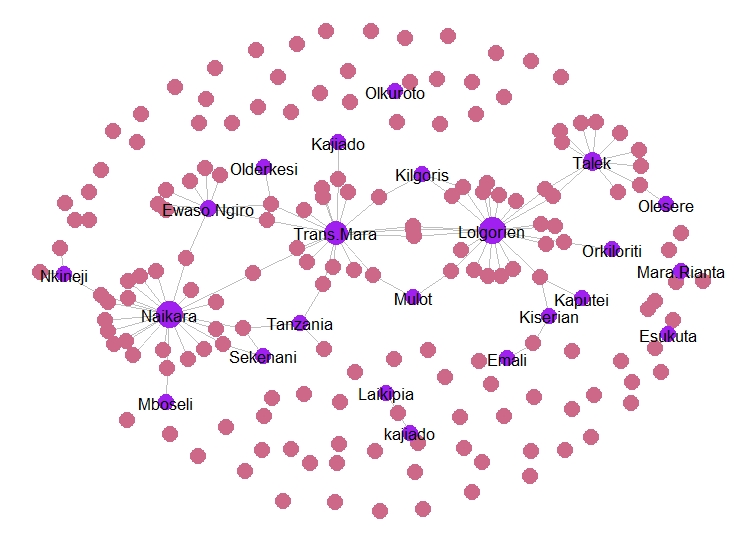

**Figure 6:** Bipartite network of selling locations following targeted removal of top five villages. Purple nodes are villages, while pale violet nodes are households.

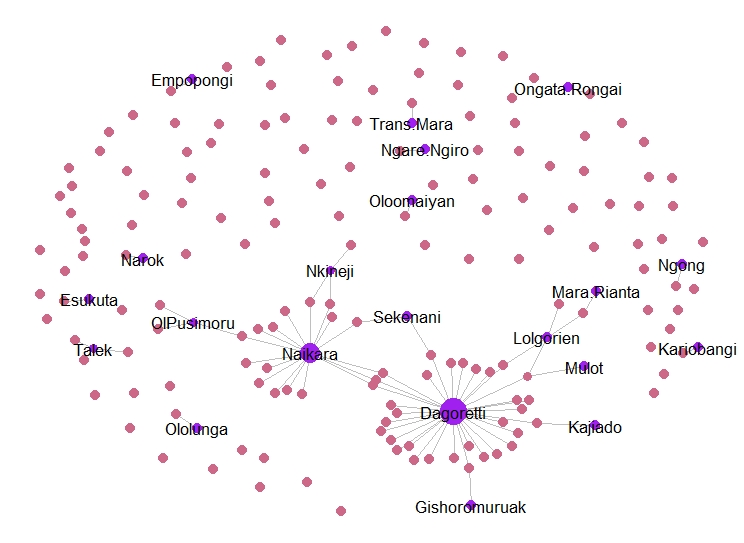
